# TypeTE: a tool to genotype mobile element insertions from whole genome resequencing data

**DOI:** 10.1101/791665

**Authors:** Clement Goubert, Jainy Thomas, Lindsay M. Payer, Jeffrey M. Kidd, Julie Feusier, W. Scott Watkins, Kathleen H. Burns, Lynn B. Jorde, Cedric Feschotte

**Author notes:** To whom correspondence should be addressed; Fax: +1; Correspondence may also be addressed to; Fax: (+1). Co-senior authors. The authors wish it to be known that, in their opinion, the first 2 authors should be regarded as joint first authors.

## Abstract

*Alu* retrotransposons account for more than 10% of the human genome, and insertions of these elements create structural variants segregating in human populations. Such polymorphic *Alu* are powerful markers to understand population structure, and they represent variants that can greatly impact genome function, including gene expression. Accurate genotyping of *Alu* and other mobile elements has been challenging. Indeed, we found that *Alu* genotypes previously called for the 1000 Genomes Project are sometimes erroneous, which poses significant problems for phasing these insertions with other variants that comprise the haplotype. To ameliorate this issue, we introduce a new pipeline -- TypeTE -- which genotypes *Alu* insertions from whole-genome sequencing data. Starting from a list of polymorphic *Alu*s, TypeTE identifies the hallmarks (poly-A tail and target site duplication) and orientation of *Alu* insertions using local re-assembly to reconstruct presence and absence alleles. Genotype likelihoods are then computed after re-mapping sequencing reads to the reconstructed alleles. Using a ‘gold standard’ set of PCR-based genotyping of >200 loci, we show that TypeTE improves genotype accuracy from 83% to 92% in the 1000 Genomes dataset. TypeTE can be readily adapted to other retrotransposon families and brings a valuable toolbox addition for population genomics.

## INTRODUCTION

Mobile element insertions (MEIs) are ubiquitous and are major contributors to genomic variation between and within species (Kidwell and Lisch 1997; Sudmant et al. 2015; Underwood, Henderson, and Martienssen 2017). Active ME families continuously generate new MEIs which segregate among individuals. Individual MEI generate structural variants (SV) between genomes (typically insertions) and can lead to complex chromosomal rearrangements through non-homologous recombination between copies (Jurka et al. 2004; Song and Boissinot 2007; Xing et al. 2009; Thomas, Perron, and Feschotte 2018). Both processes represent a substantial source of genomic instability, which has been implicated in more than 100 human genetic diseases (Hancks and Kazazian 2016), and they are also fodder for the emergence of adaptive genetic novelties (Oliver, McComb, and Greene 2013; Chuong, Elde, and Feschotte 2017; Wallace et al. 2018; Horváth, Merenciano, and González 2017; Jangam, Feschotte, and Betrán 2017)(Oliver et al. 2013; Chuong et al. 2017; Horváth et al. 2017; Jangam et al. 2017).

In humans, recently mobilized transposable elements (TEs) include members of the LINE-1, *Alu,* SVA, and a few human endogenous retroviruses (HERVs) families. Together these elements make up to >30% of the human genome, but relatively few remain polymorphic, *i.e.* being either present or absent between two genomes (Mills et al. 2007; Hancks and Kazazian 2012). Such polymorphic MEIs (pMEIs) account for hundreds to thousands of loci per individual (Stewart et al. 2011; Hancks and Kazazian 2012; Sudmant et al. 2015). The extent of pMEIs segregating in the human population is yet to be determined, but *Alu* is known to be the most common source of human pMEIs. Thus far, a little less than 20,000 *Alu* copies have been identified as segregating among 2,504 humans sampled as part of the 1000 Genomes Project (Sudmant et al, 2015; 1000 GP, Gardner et al., 2017).

*Alu* elements are powerful markers for genetic and evolutionary studies of human populations. As non-autonomous retrotransposons, *Alu*s amplify through a copy-and-paste mechanism utilizing LINE-1 machinery (Dewannieux, Esnault, and Heidmann 2003) and are inherently incapable of precise excision, providing identical-by-descent loci virtually free of homoplasy (Doronina et al. 2019). Accordingly, *Alu* have been shown to effectively track human population history (Watkins et al. 2003; Jurka, Bao, and Kojima 2011; Stewart et al. 2011; Rishishwar, Tellez Villa, and Jordan 2015). Like most MEIs, *Alu* insertions in humans are usually thought of as neutral variants that achieve fixation in the population through genetic drift (Boissinot et al. 2006; Cordaux et al. 2006). Nevertheless, more than 70 *de novo Alu* insertions are known to cause genetic diseases (Hancks and Kazazian 2016), including neurological disorders (Larsen et al. 2018; Hueso et al., n.d.). Furthermore, polymorphic *Alu* insertions have been identified as candidate causative variants in common polygenic diseases (Payer et al. 2017), and a handful have been shown to alter mRNA splicing (Payer et al. 2019). Finally, worldwide reference pMEI datasets such as those produced by 1000 GP (Sudmant et al. 2015) can be used in conjunction with gene expression data (e.g. RNA-seq) to identify loci associated with changes in gene expression (S. Wang et al. 2016).

Together these studies suggest that pMEIs, and *Alus* in particular, play an important, yet still underappreciated role in human phenotypic variation.

Recognizing the abundance and biological significance of MEIs, a growing number of software packages have been developed in the past few years to detect and map pMEIs in whole-genome resequencing (WGS) data relative to a reference genome (Goerner-Potvin and Bourque 2018). For studies of human pMEIs, Tea (Lee et al. 2012), Retroseq (Keane, Wong, and Adams 2013), Mobster (Thung et al. 2014), Tlex2 (Fiston-Lavier et al. 2015), RelocaTE2 (J. Chen et al. 2017), STEAK (Santander et al. 2017), MELT (Gardner et al. 2017), TranSurVeyor (Rajaby and Sung 2018), polyDetect (Jordan et al. 2018), and ERVcaller (X. Chen and Li 2019) are among the most recent software tools available. The algorithmic refinement dedicated to accurately detecting pMEIs, and *Alu*s in particular, in WGS data has led to an increase of the quality of the calls. Notably, the accurate detection of the presence or absence of a specific *Alu* at a precise breakpoint has improved substantially in recent years (Rishishwar, Mariño-Ramírez, and King Jordan 2016; Gardner et al. 2017; X. Chen and Li 2019).

Although the discovery of *Alu* and other pMEI alleles is generally benchmarked extensively when these methods are evaluated, far less attention has been paid to individual genotyping, *i.e.* determining whether the insertion is a homozygote or heterozygote for each individual locus. Genotyping accuracy is critical for phasing insertion polymorphisms with single nucleotide polymorphisms (SNPs) and relating insertions with expression quantitative trait loci (eQTL) and disease-risk loci identified by genome wide association studies (GWAS). Similarly, accurate genotypes are necessary to infer how the effects of drift and selection influence allele frequencies. However, genotyping accuracy of pMEI released with the 1000 GP dataset has only been estimated using 250 bp Illumina reads (accuracy estimated to 98%) (Sudmant et al., 2015). To our knowledge, only three pipelines, MELT (Gardner et al. 2017), polyDetect (Jordan et al. 2018), and ERVcaller (X. Chen and Li 2019) are maintained as tools that directly allow genotyping for non-reference pMEIs. However, MELT is the only one offering the option to directly genotype reference pMEIs (*i.e*. polymorphic elements that are annotated in the reference genome but still segregating in the population). None of these tools have been subject to a comprehensive evaluation of their genotyping performance. Given the ever-growing number of resequencing efforts, there is a pressing need to develop highly accurate genotyping tools to complement the diverse methods already available to detect the presence or absence of pMEI.

To address these issues, we have developed a new bioinformatics pipeline, TypeTE, which improves the genotyping of pMEIs located by other tools using whole genome resequencing data. Our method is based on the accurate recreation of both the presence and absence of pMEI alleles before the remapping of reads for genotyping. We benchmarked TypeTE with both low- and high-coverage data [(1000 GP phase 3 (Sudmant et al. 2015) and Simons Genome Diversity Project, (SGDP) (Mallick et al. 2016) respectively] and show, based on a collection of more than 200 PCR-based genotyping assays, that our method significantly improves genotype quality. In addition, we applied TypeTE to all polymorphic *Alu* insertions discovered in 445 human samples both present in the 1000 GP phase 3 (low-coverage WGS) and the Genetic European Variation in Disease Consortium (GEUVADIS; RNA sequencing) (Lappalainen et al. 2013). We thus provide a new high-quality genomic resource dedicated to the functional and evolutionary analysis of polymorphic *Alu* insertions.

## MATERIAL AND METHODS

### Pipeline implementation

#### Non-reference MEI

TypeTE-*non-reference* is designed to genotype insertions not found in the reference genome (Figure 1A, Supplementary Figure S1). Based on the information provided in a vcf file (such as produced by MELT), the location and orientation of each *Alu* insertion are first collected. For each breakpoint, reads that are mapped in a window of 500 bp (250 bp upstream and downstream of the breakpoint) are extracted. The mates of discordant reads (mapping somewhere else in the genome) are also extracted from the BAM file of each individual. The reads from all individuals for each locus are then combined, and a local *de-novo* assembly of all the reads is attempted using SPAdes v3.11.1 (Bankevich et al. 2012). Minia (v2.0.7) (Chikhi and Rizk 2013) is used an alternate assembler when SPAdes failed to generate an assembly of the sequences (‘scaffolds.fasta’). The genomic locations where mates of discordant reads are mapped are identified and intersected with the respective RepeatMasker track (we used the coordinates version hg19 for 1000 GP data and hg38 for the SGDP data; Repbase version 20140131). Using a majority rule, the most likely *Alu* subfamily consensus for the copy inserted at that locus is identified. To verify orientation and identify target site duplications (TSDs), homology-based searches are performed. First, the identified *Alu* consensus is searched with blastn (v. 2.6.0+) against the assembled contigs. Then, a second blastn is performed using the genome reference sequence (500 bps window) against the assembled contigs. The contig with the highest score from the query *Alu* and the reference sequence is selected and searched for target site duplications flanking the MEI. To identify the strand of the MEI, the sequence flanking the insertion in the contig is further compared with the reference sequence. For each MEI, the two alleles are reconstructed as follows: a new window of +/− 500bp is extracted upstream and downstream of the breakpoint predicted by MELT. This represents the “absence” allele. To recreate the “presence” allele, TypeTE first removes the predicted TSDs from the extracted reference sequence and inserts the fully assembled MEI with its two TSDs in the correct orientation. If the assembly fails to generate a complete sequence of the MEI with flanking TSDs, the TSD predicted by the TE detection program (in our case MELT) is duplicated and placed at the 5’ and 3’ end of the consensus MEI in the composite allele.

**Figure 1:**
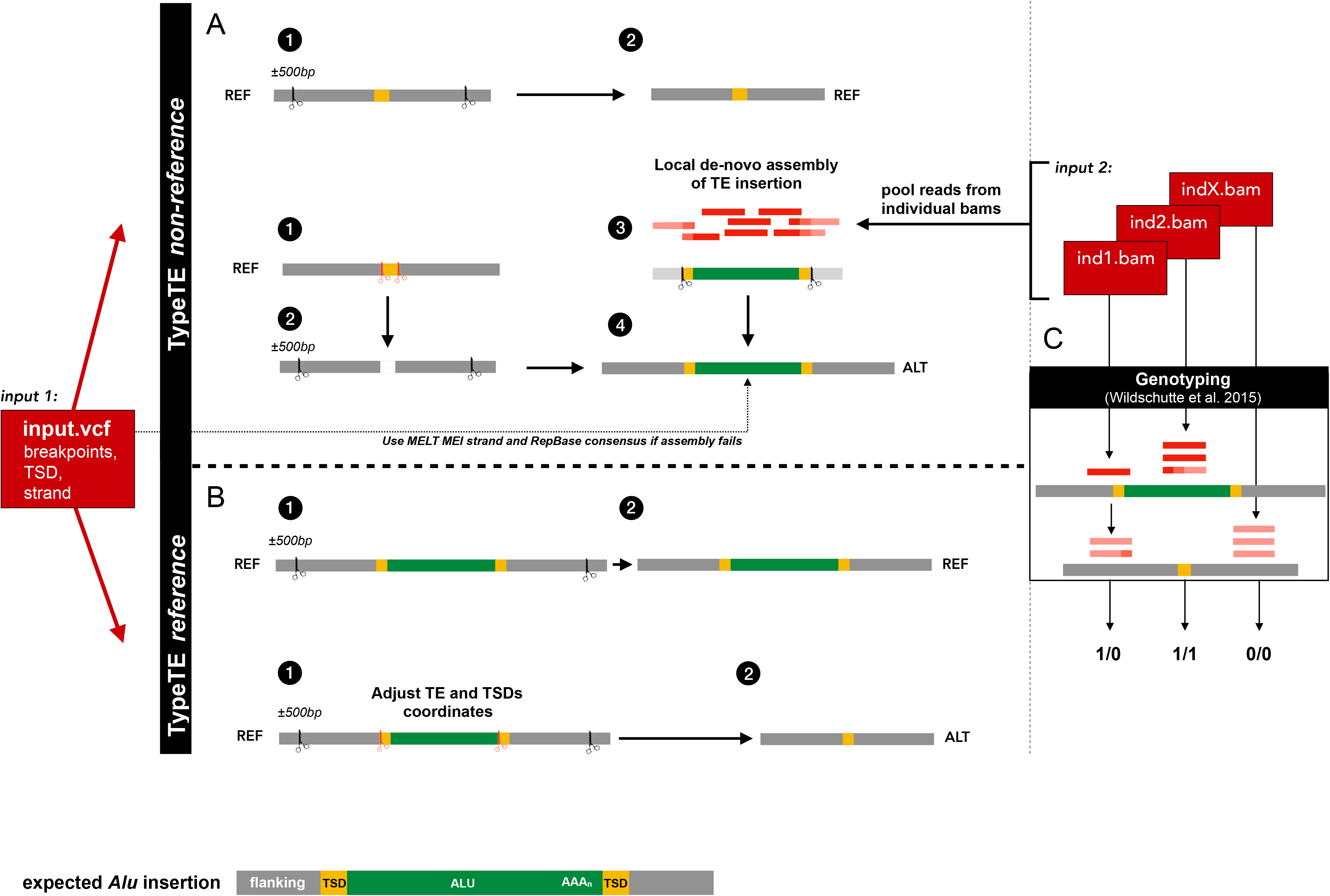
Overview of the TypeTE pipeline. TypeTE is divided in two main scripts. The first (A) genotypes non-reference insertion (TypeTE-nonref) and the second (B) genotypes reference pMEI (TypeTE-ref). (A) TypeTE-ref creates the reference allele (REF) by extracting +/−500 bps from the *Alu* predicted breakpoint. The alternate allele (ALT), corresponding to the pMEI presence is made by 1-2) removing the predicted TSD from the +/−500 bps extracted sequence. Then, for each locus, read pairs (including discordant mates) are extracted from the individual bam files and are pooled for local assembly (3). If TSDs are identified in the assembly, the sequence is then inserted onto the flanking (4). In case the assembly is incomplete, the Repbase consensus for the predicted TE family is inserted instead (4). (B) The REF allele is created after extraction of +/−500 bps from the 5’ and 3’ ends of the adjusted *Alu* position (including TSDs). The ALT allele is then created removing the *Alu* sequence and 1 TSD from the same extracted sequence. (C) Genotyping. For each locus, read-pairs of each sample are extracted in a 500 bps window centered on the predicted breakpoint. For each sample, these reads are then mapped to the two alleles and genotype likelihood are computed.

#### Reference TE

*TypeTE-reference* determines genotypes of *Alu*s in the reference genome that are polymorphic in other individuals (Figure 1B, Supplementary Figure S2). In this case, no reads are extracted from the original alignments to reconstruct the alternate allele. However, the exact coordinates and TSDs of each MEI in the reference genome are reassessed as follows: the breakpoints identified from MELT for the location of the reference TE are further refined using the corresponding RepeatMasker annotation track to identify the exact location and orientation of each TE inserted in the reference genome. At first, the longest reference *Alu* elements that are within +/−50 bps of the predicted MELT breakpoints are extracted. If none is found within that boundary, *Alu*s within +/−110 bps of the predicted breakpoints are collected. However, we did not find any difference in the number of elements identified after increasing the boundary up to 200bps. The flanking sequence of the TE sequence is also extracted and TSDs and their coordinates are identified whenever possible. Then, based on these new coordinates, a region of +/−500 bps upstream and downstream of the 5’ and 3’ end of the MEI is extracted from the reference genome. This constitutes the “presence” allele. The “absence” allele is defined by removing the TE sequence and one TSD from the reference genome.

#### Genotyping

TypeTE automatically generates input files and parallelizes the method developed by Wildschutte et al. (Wildschutte et al. 2015), called *insertion-genotype*, to genotype each *Alu* insertion in every individual. Briefly, read-pairs with at least one read mapping to the target locus are extracted and mapped against the reconstructed insertion and empty site alleles using bwa (v. 0.7.16a) (H. Li and Durbin 2009). The number of reads that align to each allele, and their associated mapping quality values are tabulated and likelihoods for the three possible genotype states are calculated (H. Li 2011). Reads that map equally well to the empty and insertion alleles are assigned a mapping quality of 0 by bwa (H. Li and Durbin 2009) and do not contribute to this calculation. Additionally, read pairs are required to partially align to the repeat sequence and pairs that align entirely within the target repeat sequence are ignored, since these reads may not be specific to the targeted locus. By default, the genotype with the highest likelihood is chosen, but the resulting likelihoods may optionally be used as inputs to downstream programs which estimate genotypes based on patterns across multiple samples and sites. After genotyping, individual per-sample VCFs are concatenated.

### Evaluation of the 1000 GP genotypes quality and TypeTE performance

#### Genotype calling

In order to evaluate the quality of the *Alu* genotype calls available in the 1000 GP phase 3 structural variants (SV) dataset ([Sudmant et al. 2015], average depth of coverage 7.4X), we gathered the genotypes available for both non-reference (indicated by “<INS:ME:ALU>“ in the available VCF file) and reference (tagged with “SV_TYPE=DEL_ALU”). We ran TypeTE-reference and TypeTE-non-reference on the same loci as well as MELT-discovery (non-reference) and MELT-deletion (reference) using its version 2.1.4 (referred to as MELT2 for the remainder of the manuscript) in order to take into account, the most recent changes added to its genotyping module. Additionally, we tested the performances of TypeTE with samples from the SGDP (Mallick et al. 2016), which has higher coverage (average 42X).

In the 1000 GP data, we ran TypeTE and MELT2 on 445 CEU, TSI, GBR, FIN and YRI individuals, also present in the Geuvadis dataset (RNA-seq) (Lappalainen et al. 2013). In the 1000 GP dataset released by Sudmant et al. (Sudmant et al. 2015), Alu genotypes were produced by MELT (first version) for non-reference insertions. However, polymorphic reference Alu insertions were first discovered along with other genomic deletions with a set of SV detection tools (BreakDancer, Delly, CNVnator, GenomeSTRiP, Variation-Hunter, SSF and Pindel), then genotyped with the same algorithm as any other SV (Sudmant et al. 2015). Because the sample size we used was smaller than the original one (n = 445 vs n = 2504), MELT2 did not recover all loci genotyped by the 1000 GP and TypeTE. Also, probably because of changes in the newer version, some Alu breakpoints were slightly different between the Sudmant et al. (Sudmant et al. 2015) dataset and the MELT2 output. Thus, in order to reconcile and compare the three datasets, bedtools intersect (v.1.5) (Quinlan et al. 2010) was used with a window of +/− 30bp around each original 1000 GP Alu breakpoint. Finally, the predicted genotypes were compared to PCR assays of 108 non-reference and 43 reference loci in 42 individuals from the CEU population (see next section).

For the SGDP data, reference and non-reference polymorphic *Alu* insertions were called using MELT2 in 14 publicly available individuals from the South Asian population for which we had access to DNA. The genotypes of the loci discovered were then determined using TypeTE and compared to 9 non-reference and 67 reference loci previously genotyped by PCR in these 14 samples (Watkins et al. 2003).

#### PCR typing in a subset of 1KGP and SGDP dataset

Non-reference (*108*) and reference (*43*) *Alu* loci identified in 1000 GP were tested in a 30-trio reference panel of CEPH CEU individuals (42 individuals were evaluated by PCR and sequenced in 1000 GP) (HAPMAPPT01, Coriell Institute for Medical Research). Primers flanking the *Alu* insertion site were selected using Primer3 (Untergasser et al. 2012). PCR amplifications were performed using *OneTaq* Hot Start Quick-Load 2x Master Mix (New England BioLabs) using 3-step PCR (initial denaturation: 94°C, 15”, (94°C, 15’’; 57°C, 15’’; 68°C, 30’’) for 30 cycles; final extension 68·C, 5’). Sequences for 20 new primer pairs are available in Table S1; the remainder are available in (Payer et al. 2017). Accuracy was evaluated by replication in duplicate samples and by evaluating the number of Mendelian errors in related individuals. Non-reference (9) and reference (67) *Alu* loci were previously genotyped by PCR in 14 South Asian samples present in the SGDP dataset (Watkins et al. 2003). Primers around each *Alu* insertion were selected using Primer3 (Untergasser et al. 2012). PCR amplification was performed using three-step PCR (initial denaturation: 94°C, 3’; (94°C, 15’’; 60°C, 15’’; 72°C, 30’’) for 30 cycles; final extension 72·C, 5’) in 1X PCR buffer (10mM Tris, pH 8.3, 50mM KCl, 1.5 mM MgCl2) with 200 uM dNTPs, 10 pmol each primer, and 1U Taq polymerase. Annealing temperature was adjusted for each primer set. DMSO (5-10%) was used to improve amplification for some loci.

### Effect of genotype corrections on the *Alu* insertion discovery

In some cases, new genotyping changed the presence/absence status of an *Alu* insertion for a given genome. We define a false positive (FP) as a case in which an *Alu* copy is called present, either homozygote or heterozygote in one sample, while the PCR reported it absent. A false negative (FN) is recorded when an *Alu* is called absent (homozygote absent) while it is called as either homozygote present or heterozygous by PCR. True positive (TP) and true negative (TN) are the same calls (presence/absence), respectively, being validated by PCR. For each dataset and method, we calculated the sensitivity (ability of the method to discover a MEI: TP/(TP+FN)), the precision (or positive predictive value: TP/(TP+FP)) as well as the F1 score as described by Rishishwar et al (Rishishwar, Mariño-Ramírez, and King Jordan 2016), which corresponds to the harmonic mean of sensitivity and precision and summarizes the overall performance of each method.

### Estimation of mappability scores

The mappability scores are downloaded for the GRCh37/hg19 version of reference assembly for 100mers (ftp://hgdownload.soe.ucsc.edu/gbdb/hg19/bbi/wgEncodeCrgMapabilityAlign100mer.bw). The downloaded file is processed (Kent et al. 2010) and is converted to bed format (Neph et al. 2012). These data are stored in an indexed mysql table. The mappability scores for genomic regions in the flanking region (+/−250bps) of the predicted *Alu* breakpoint for non-reference insertions and flanking region (+/−250bps) of the reference *Alu* insertions are extracted from the table, and the mean of the mappability scores is recorded in a dedicated table and is provided with the output files.

### Calculation of local read depth

The average read depth at genomic regions in the flanking region (+/−250bps) of the predicted *Alu* breakpoint for non-reference insertions and flanking region (+/−250bps) of the reference *Alu* insertions is calculated using samtools (Version: 1.4.1). Only reads with a mapping quality of 20 or more (mapped with > 99% probability) and bases with a quality of 20 or more (base call accuracy of > 99%) are counted.

### Inbreeding Coefficient (*Fis*) estimates

In order to assess how genotype quality affects common population genetics summary statistics, we computed the per locus inbreeding coefficient (*Fis*) for the loci assayed by PCR. *F*_is_ is a common metric used in population genetics to assess the excess (*F*_is_ < 0) or the depletion (*F*_is_ > 0) in heterozygotes relative to the expected genotypes proportion at Hardy-Weinberg equilibrium. Allele frequencies were calculated using the genotypes produced by each method (1000 GP, MELT2, TypeTE and PCR) as follows:

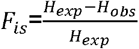

with H_exp_ = 2pq, p = presence allele insertion frequency, q = (1-p) H_obs_ is the observed number of heterozygotes.

All statistical analyses were carried out with R version 3.5.1 (R Core Team 2018).

## RESULTS

### Concordance of the 1000 GP dataset genotypes with PCR assays

*Alu* genotype predictions in the 1000 GP phase 3 release (Sudmant et al. 2015) were called using MELTv1.0 (first version) for non-reference loci and a combination of SV tools (Sudmant et al. 2015) for reference insertions. To assess their accuracy, we compared them to an assembled collection of 108 non-reference and 43 reference loci genotyped by PCR (Figure 2A and Table 1) in 42 individuals (see methods). To ensure accuracy in genotyping validations, PCR assays were performed using all (30) trios of the CEPH CEU, and in all cases, no Mendelian errors in the transmission of alleles from parents to offspring were seen (see methods). Presence of both “empty” and “filled” alleles (with and without *Alu*) were confirmed by the presence of bands of expected size in the agarose gel electrophoresis and in most cases with Sanger sequencing. Upon comparing the genotype predictions to PCR assays, we found that the 1000 GP phase 3 release had an overall concordance rate with the PCR (total number of prediction identical to the PCR generated genotypes/ total number of predictions) of 83.31% (3649/4380) for non-reference *Alu* insertions and 80.72% (1248/1590) for reference insertions.

**Figure 2:**
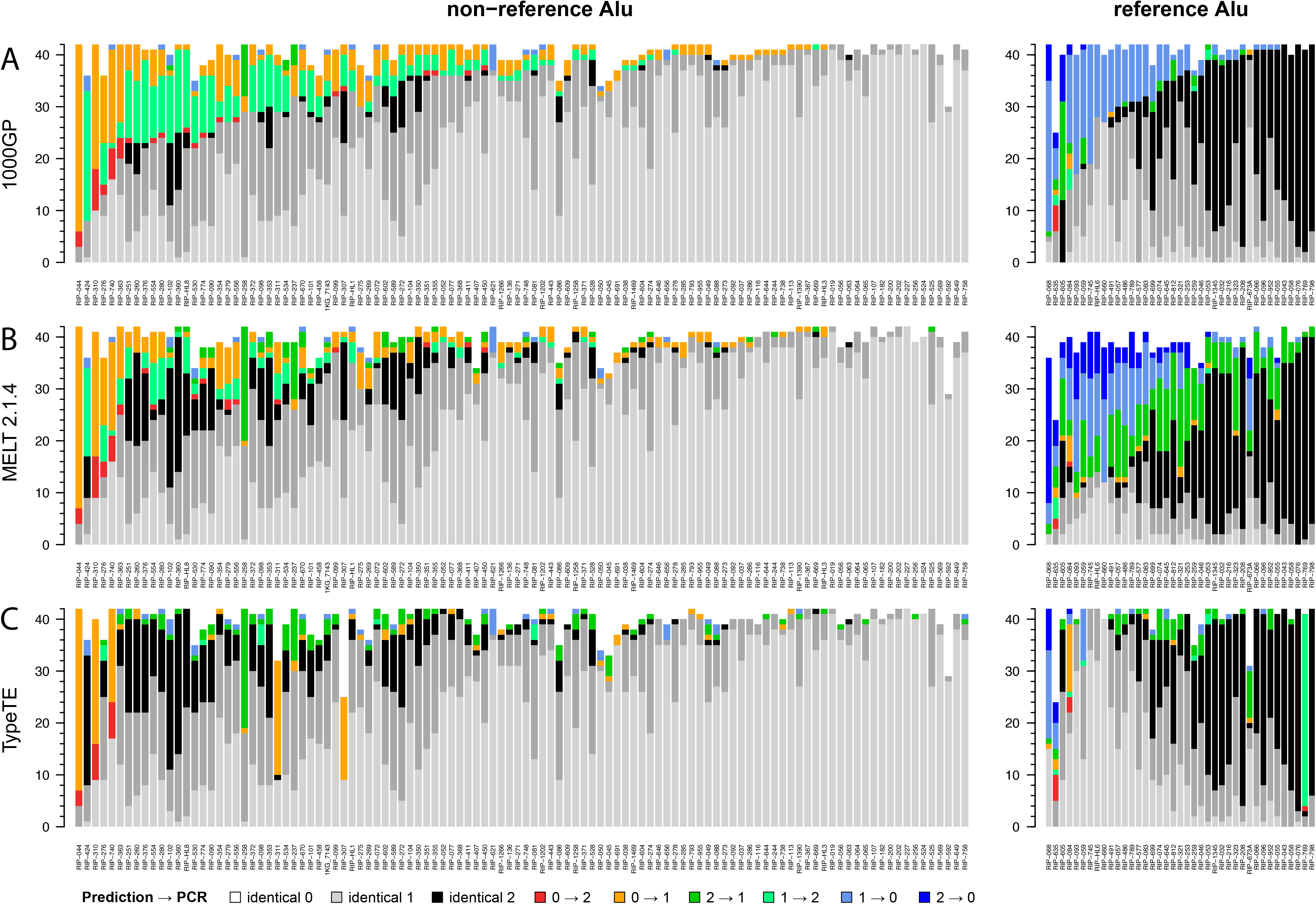
Comparison of the predicted genotypes in the 1000 GP dataset with PCR-assays in 42 CEU individuals. Each vertical bar represents one locus, and match or error regarding the genotype for each individual are piled up on the Y axis and color coded according to the legend. NA values (no genotype predicted or failed PCR) are removed from the plot. >

**Table 1.**
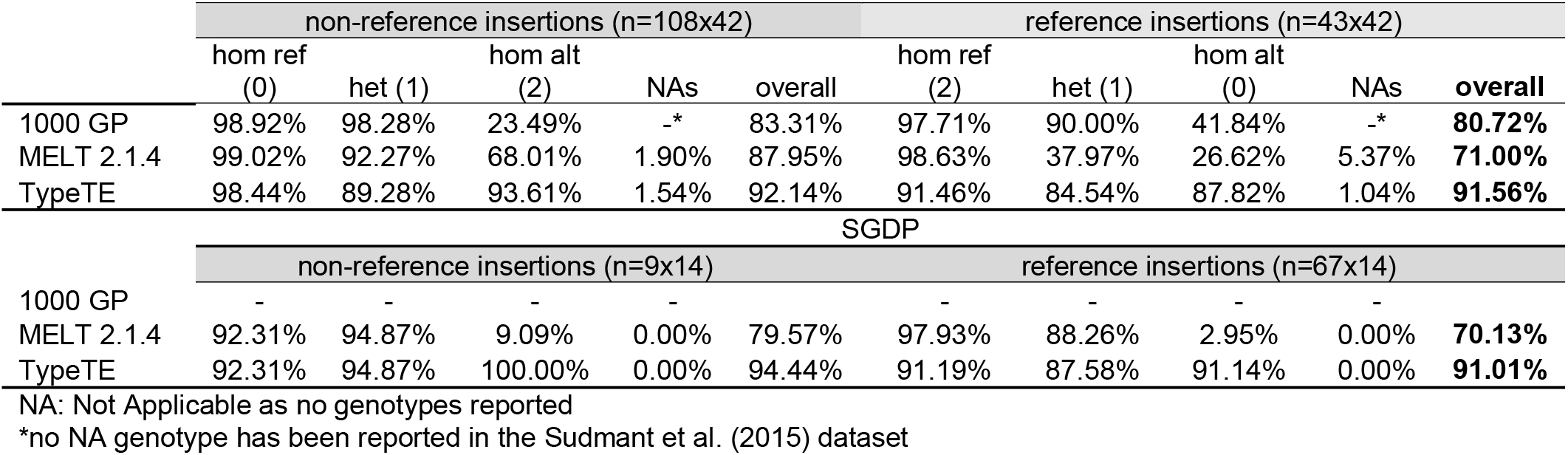
Genotype prediction accuracy (%) for each category of insertions when compared with PCR generated genotypes

### TypeTE pipeline overview

In order to improve the quality of *Alu* genotyping by short read sequencing analysis, we developed TypeTE which allows the re-genotyping of both reference and non-reference *Alu* insertions. The pipeline is divided into two main modules: the *non-reference* module genotypes *Alu* insertions absent from the reference genome, while the *reference* module genotypes *Alu* insertions present in the reference genome. Details about the implementation of each module are given in the Material and Methods section (Figure 1,Supplemental Figure S1 and Supplemental Figure S2). The basic principle of TypeTE is to recreate the most accurate sequences for the two alleles of each insertion (presence and absence). TypeTE currently uses a VCF file such as produced by a TE discovery tool such as MELT to locate each individual TE insertion. The pipeline then performs an independent analysis of each predicted locus and collects the information regarding the insertion. After allele reconstruction (see Material and Methods), the individual reads mapping to each insertion locus are extracted from the original alignment file (bam) and mapped against the reconstructed alleles for genotyping using an automated and parallelized version of the method developed by Wildschutte et al. (2015). A new VCF file with the corrected genotypes and genotypes likelihoods is then produced.

### TypeTE performances

In order to assess the accuracy of the predictions made by TypeTE, we ran the pipeline on a subset of 445 individuals of European and African ancestry included in the 1000 GP dataset (see Material and Methods). These samples were selected because they are both represented in the 1000 GP (WGS) and GEUVADIS (RNA-seq) datasets, allowing functional analyses of pMEIs. We also compared the performance of TypeTE with a recent version of MELT (version 2.1.4, abbreviated as MELT2) using the packages MELT-discovery and MELT-deletion on the same sample. TypeTE and MELT2 genotypes were then compared to 108 non-reference and 43 reference insertion for which we have collected or generated PCR genotypes. With non-reference insertions, MELT2 shows increased concordance with the PCR compared to the original 1000 GP calls, with 87.95% (vs 83.31%; +131/4298 accurate genotypes) of the predicted genotypes matching the experimental results. TypeTE further increases the concordance of the genotype prediction, achieving a rate of 92.14% (+325/4313 accurate genotypes compared to original 1000 GP release). For reference insertions, MELT2 had a lower concordance than the original 1000 GP predictions, with only 71% (vs. 80.72%; −374/1504 genotypes) of the genotypes matching the PCR results, while TypeTE achieved 91.56% concordance (+ 141/1575 genotypes). Note that the total number of genotypes considered correspond to the total number of predictions available and doesn’t take into account here the missing genotypes.

We further tested the genotyping performance of MELT2 and TypeTE with the SGDP (Mallick et al. 2016) data, which benefits from a higher depth of coverage than the 1000 GP data (42x vs 7.4x). We tested the concordance of the predicted genotypes with 67 reference and 9 non-reference *Alu* loci in 14 individuals previously genotyped by PCR (Watkins et al. 2003). MELT2 has a concordance rate of 70.13% for reference loci while TypeTE matches the PCR results for 91.01% of the predicted genotypes (+181 correct genotypes; Figure 3 and Table 1). Finally, for the 9 non-reference loci that were experimentally genotyped, the concordance rate is 78.57% for MELT2 and 94.44% for TypeTE (+ 20 correct genotypes).

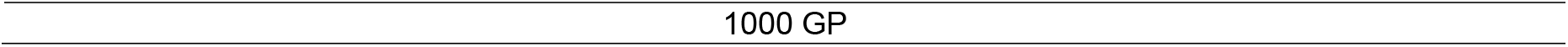

**Figure 3:**
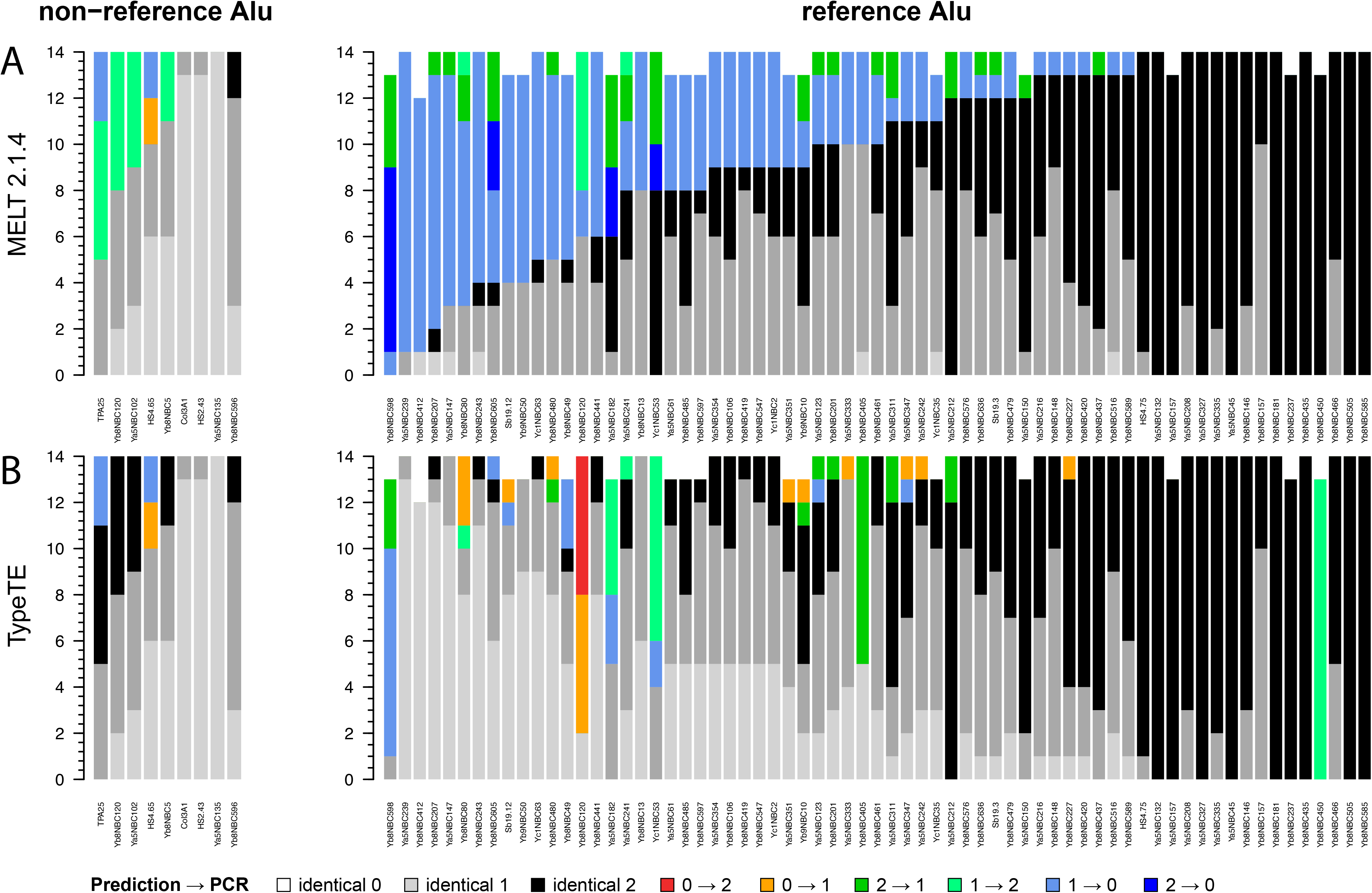
Comparison of the predicted genotypes in the SGDP dataset with PCR-assays in 14 South Asian individuals. Each vertical bar represents one locus, and match or error regarding the genotype for each individual are piled up on the Y axis and color coded according to the legend. NA values (no genotype predicted or failed PCR) are removed from the plot. >

In order to analyze in detail, the genotyping performances of each method, we calculated the concordance rate by genotype category (0 or (0/0): homozygote absent, 1 or (0/1): heterozygote, 2 or (1/1): homozygote present) corresponding to the percent of correct genotypes in one category to the total number of calls for this category (Table 1). Additionally, we report the percentage of unascertained loci (NA genotypes) for each method.

We then investigated how the concordance between predicted and PCR genotypes is distributed across loci and individuals by calculating the average concordance rate at each locus (total number of correct genotypes at a locus / total number of individuals with a predicted genotype). Regardless of the genotype category (reference / non-reference), TypeTE has a higher average concordance rate per locus, as well as lower variance for this value, than the other methods (Figure 4). The greatest improvement was when the genotypes of reference insertions were compared to MELT2, where the concordance rate of TypeTE is always significantly higher (Tukey’s HSD, *P* < 0.05).

**Figure 4:**
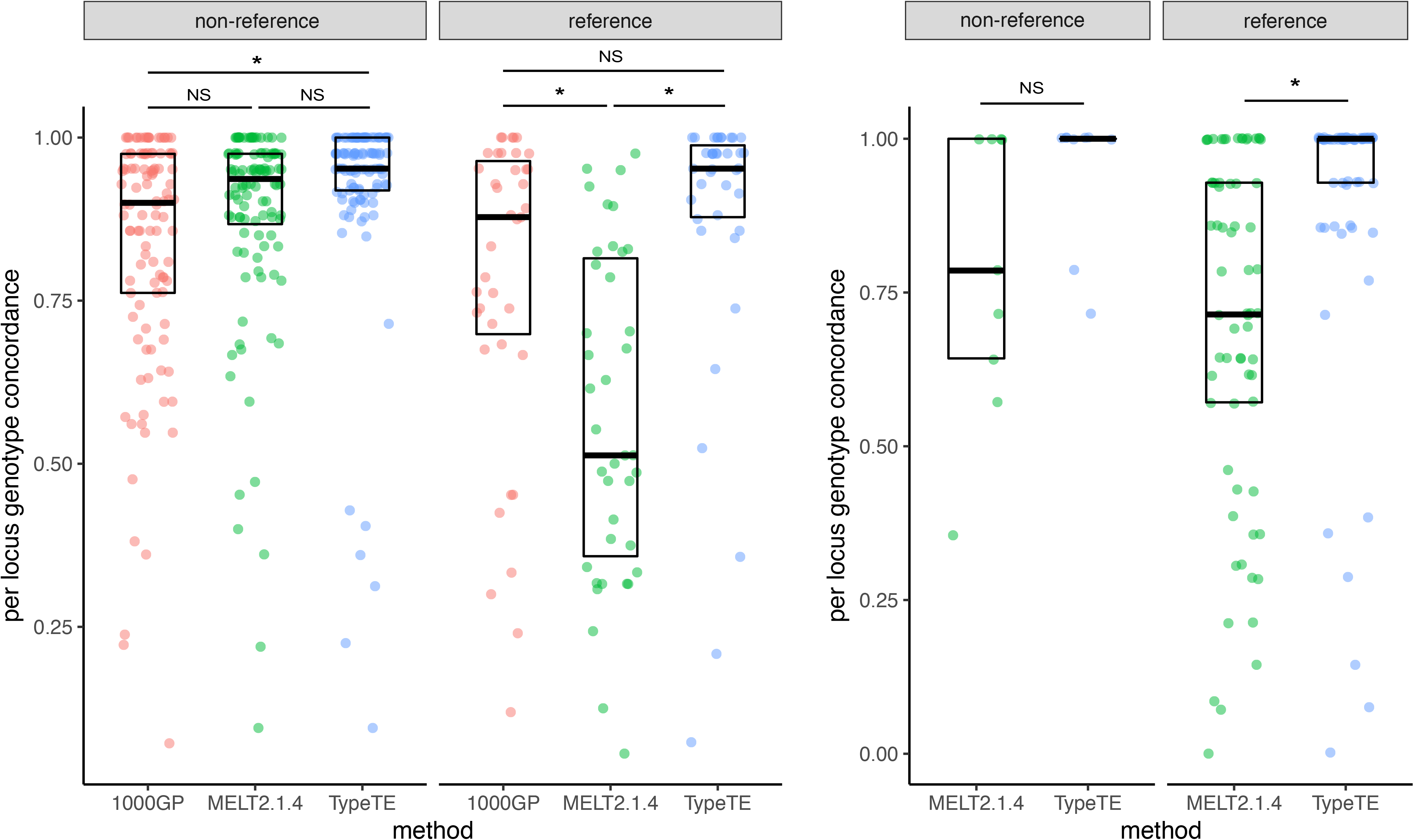
Average error rate per locus across methods and datasets. Different letters indicate significant difference. Tukey’s HSD, *P* < 0.05; NS: Not significant>

For each locus assayed by PCR in the 1000 GP dataset, we also examined whether the mappability and local read coverage affect genotyping predictions for TypeTE. We do not find a significant correlation between genotype concordance and the mappability score (0.1 - 1) computed in a 500-bp window around the MEI breakpoints (Supplementary Figure S3. Pearson’s product-moment correlation, r = 0.20, *P* = 0.281 for non-reference loci and r = 0.13, *P* = 0.414 for reference loci). We also found that the average depth of coverage for a given locus (4.69 X – 10.01 X) is not correlated to genotyping concordance for both reference (r = 0.12, *P* = 0.4538) and non-reference insertions (r = −0.01, *P* = 0.957) (Supplementary Figure S4). We conclude that at least for the loci tested by PCR, the level of repetitiveness of the flanking sequence of individual *Alu* insertions and the local read depth do not appear to influence the genotyping performance of TypeTE.

### Effect of genotype corrections on variant discovery

Different methods can assign different genotypes for some loci due to the inherent differences in their approach or due to locus specific features. For example, a heterozygous locus for the presence of *Alu* can be genotyped either as homozygous presence or absence by different methods. We first converted the genotypes into presence/absence calls in order to assess sensitivity, precision (positive predictive value), and the overall detection accuracy, summarized by the F1 score (harmonic mean of sensitivity and precision, see Material and Methods) for each method considering PCR results as true genotypes. TypeTE received the highest F1 score in each dataset (1000 GP or SGDP) and for both types of insertion (reference or non-reference) (Fig 5). The small number of loci tested for the SGDP-non-reference dataset (n = 9) did not allow us to find significant differences between the methods; however, we show that the increased F1 score of TypeTE with the 1000 GP-non-reference loci is due to a significant increase of the sensitivity compared to the other methods. Interestingly, the higher F1 score of TypeTE with reference insertions (both for the 1000 GP and SGDP datasets) is, in these cases, due to significantly higher precision (TP/(TP+FP)).

**FIGURE 5:**
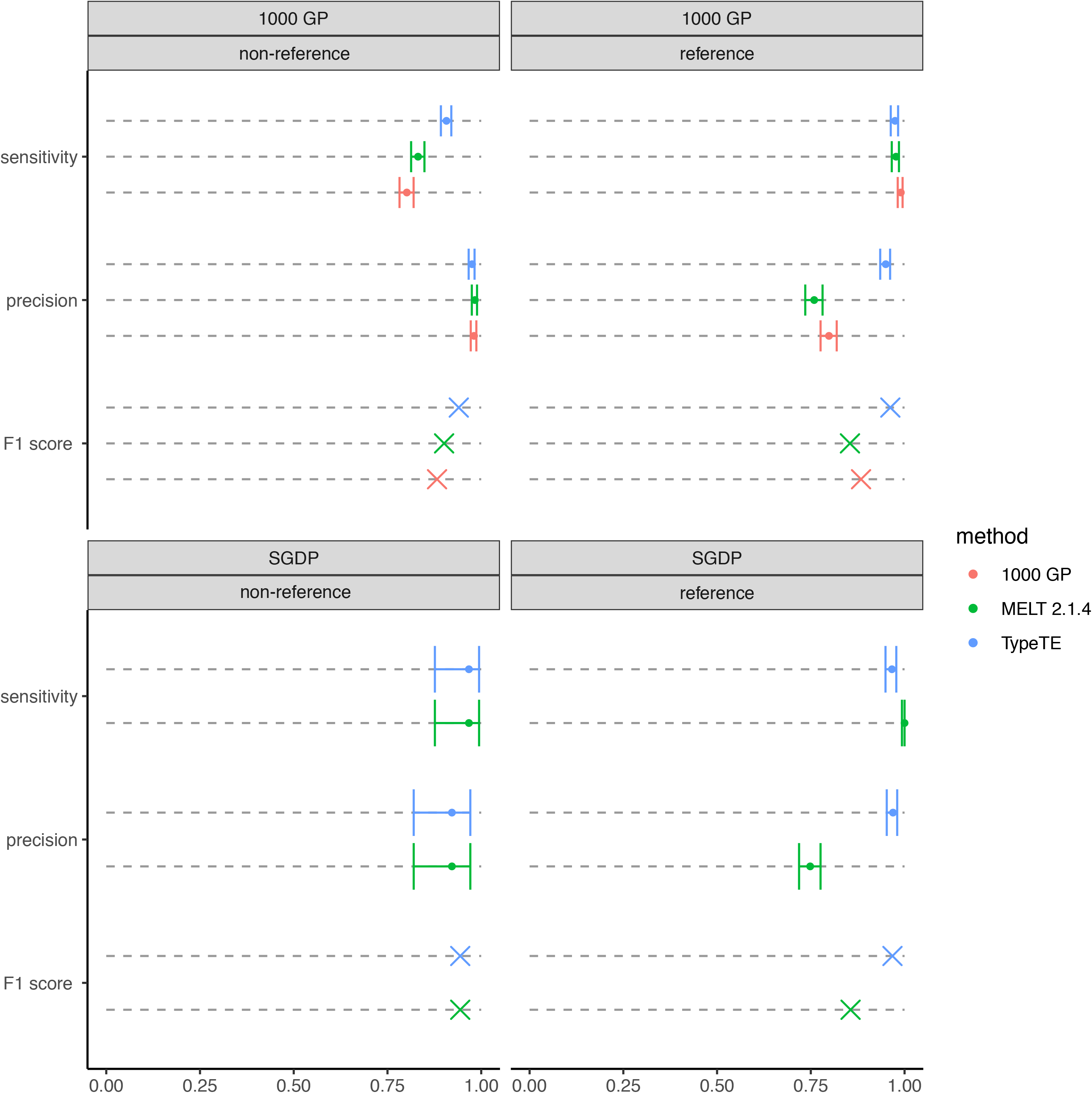
Effect of method and dataset on variant discovery performance. Sensitivity, precision and F1 score are compared for each dataset (1000 GP and SGDP) according to the type of insertion (non-reference vs reference) and the genotyping method used (1000 GP, MELT2.1.4 and TypeTE). Error bars: 95% confidence interval. Non-overlapping intervals denotes a significant difference between scores.

### Influence of re-genotyping on population genetics statistics

To illustrate the importance of accurately genotyping of *Alu*s, we calculated the population-wise inbreeding coefficient (*F*_is_) for each locus in 42 individuals of the CEU cohort (1000 GP) and 14 individuals of the South Asian cohort (SGDP). Compared to the original 1000 GP and MELT2 genotypes, the *F*_is_ values calculated with TypeTE genotypes are concordant with the ones based on PCR genotypes. These results are even more striking when only reference loci are considered: while TypeTE and PCR estimates of *F*_is_ are centered at 0, MELT2 and 1000 GP genotypes suggest a clear deviation of most loci from Hardy-Weinberg equilibrium (Figure 6). We note that estimates of the *F*_is_ are more variable using the SGDP data, which can be explained by its smaller sample size and a higher population subdivision (e.g. castes) than the 1000 GP dataset.

**FIGURE 6:**
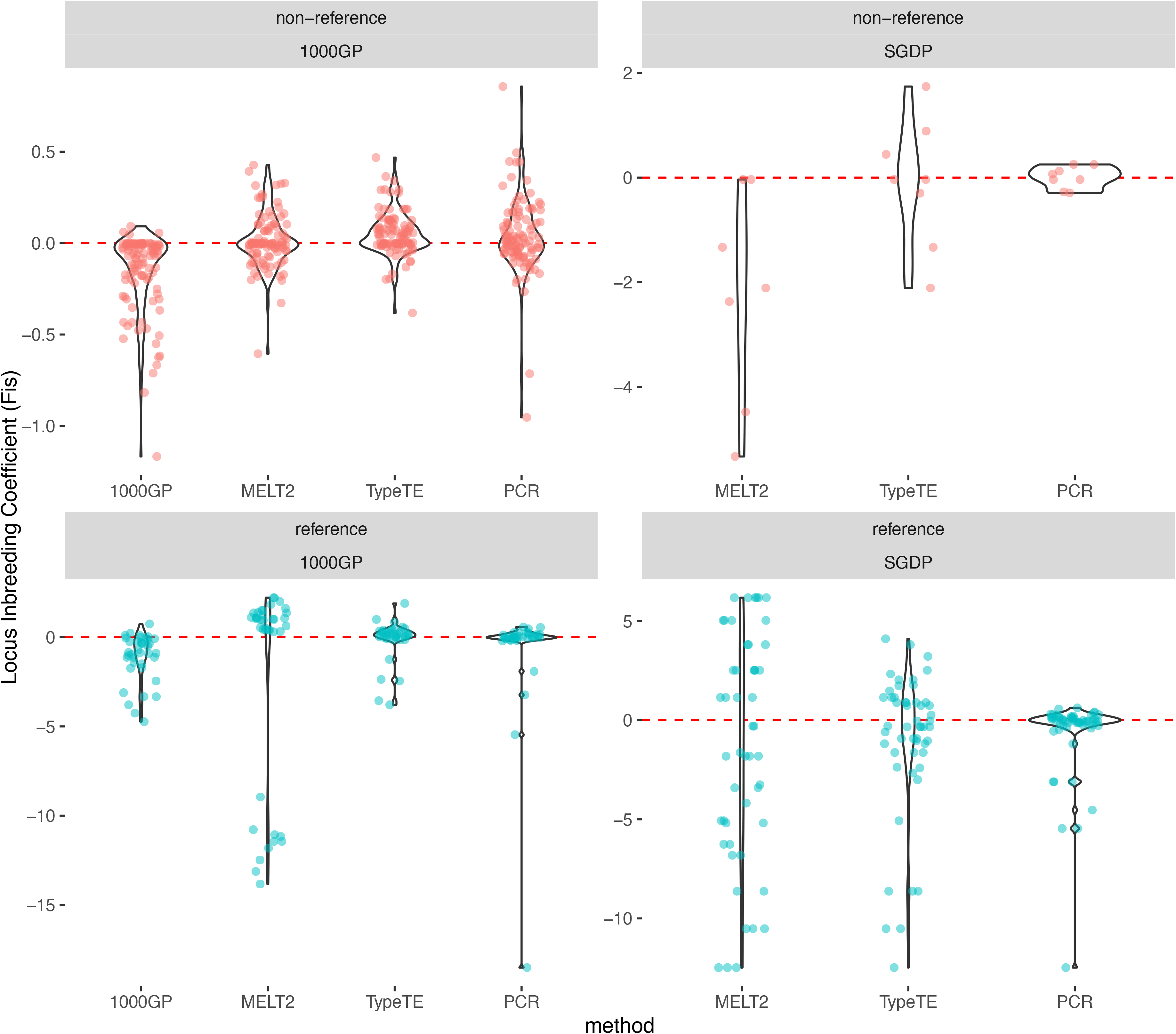
Per locus inbreeding coefficient (Fis). The Fis is estimated for each locus using the alleles frequencies given by each method (1000 GP: original 1000 GP genotypes, MELT2, TypeTE and PCR assays) and for each of the 1000 GP (n = 42 individuals) and SGDP (n = 9 individuals) datasets. Red dashed-line: expected Fis at Hardy-Weinberg equilibrium (Fis = 0).

### Influence of the dataset quality on genotype prediction

To discover factors specific to each dataset that influence genotype prediction, we compared the results obtained in the 1000 GP dataset (average depth of 7.4 X) with the results from analysis of the SGDP (average depth of 42X) to the respective PCR genotypes. The provenance of the dataset does not influence the variant discovery abilities of MELT2 and TypeTE (Supplementary Figure S5). However, we observe that the percentage of unascertained loci differs between the 1000 GP and SGDP datasets. About 1% to 5.4% of genotypes are not ascertained by either MELT2 and TypeTE in the 1000 GP dataset, probably due to low coverage. Conversely, all SGDP loci are called in every individual for the SGDP dataset (Table 1).

## DISCUSSION

The purpose of TypeTE is to provide automatic and reliable genotyping of pMEIs, especially *Alu*s from short read, whole genome or targeted interval sequencing. To our knowledge, MELT (Gardner et al. 2017) is the only tool with continued support and documentation that allows direct genotyping of both reference and non-reference pMEI. While its performance for variant discovery has made it a popular tool for pMEI mapping, to our knowledge its performance at genotyping has never been comprehensively tested. Moreover, there was no formal testing of the genotype quality concerning pMEI reported in the Phase 3 release of the 1000 GP. Thus, we thoroughly tested the *Alu* genotype predictions made for the 1000 GP, a recent version of MELT (v. 2.1.4), and TypeTE by assembling the results of more than 200 locus-specific PCR genotyping assays.

Combining reference and non-reference pMEI, our analysis indicates that 82% of the genotypes reported by the 1000 GP are consistent with the PCR assays which we consider the ‘gold standard’ and wherein accuracy was evaluated by comparing duplicate samples and verifying the absence of Mendelian errors in related individuals. This estimate of genotyping accuracy is much lower than a previous estimate of 98% based on long (250bp) Illumina reads (Sudmant et al. 2015). Genotypes reported by the 1000 GP were estimated using the first version of MELT for non-reference loci, but genotyping methods developed for other structural variation (indels, inversions, etc.) were used for reference insertions. While MELT2 appears to offer a noticeable improvement over its first version for genotyping non-reference pMEI, its overall genotyping performance is diminished when applied to reference loci, with genotyping errors reaching more than 20% in based on our PCR assays. For both categories of loci, most errors are caused by the underestimation of homozygous genotypes carrying the alternative allele relative to the reference genome (Table 1). We note that for non-reference insertions, MELT’s genotyping algorithm benefited from improvements deployed in the version tested (v2.1.4) compared to its original release, in particular to detect homozygous insertion (1/1). However, this increased sensitivity to detect pMEI alleles from read alignments seems to be accompanied by a reduced power to detect “absence” alleles for reference insertions (MELT-deletion module). Such errors are consequential for population genetics analysis because they lead to inaccurate estimation of population genetics parameters. For example, calculation of the inbreeding coefficient (*F*_is_) shows that the original release of the 1000 GP genotypes was overestimating heterozygotes, leading to negative and likely inaccurate values of *F*_is_ (Figure 6). Genotypes obtained with MELT2 improve these estimates for non-reference insertions, but the results appear less accurate when computed from a small sample and they are more inaccurate for reference insertions. These issues underscore the need for a tool dedicated to the genotyping of pMEI.

Toward this goal we developed TypeTE and applied it to genotype both reference and non-reference *Alu* insertions. Our benchmarking data show that TypeTE has an average concordance rate of 91% or greater with PCR-based genotyping. Importantly, TypeTE maintains a genotyping accuracy greater than 84% under all genotyping scenarios. While TypeTE performs better than MELT v1 (1000 GP) and MELT2 for non-reference insertions, the most significant improvement is for reference insertions. In particular, the genotypes predicted by 1000 GP and MELT2 never reached more than 41.8% concordance with the experimental results when the PCR called a homozygote absence (0/0); by contrast, TypeTE predicted these genotypes with more than 87% concordance in the two datasets tested (1000 GP and SGDP). Consequently, calculation of *F*_is_ based on TypeTE genotypes shows better concordance with that based on PCR-derived genotypes, and fits the neutral expectation as we observe no deviation from Hardy-Weinberg equilibrium for a single human population (Hosking et al., 2014).

The principal difference between TypeTE and MELT derives from characteristics of the actual data on which the genotyping is performed. While both methods implement the core genotyping algorithm described by Li (H. Li 2011), TypeTE relies on a strategy based on re-alignment of the reads against both presence and absence alleles before computation of the genotype likelihoods, an approach initially introduced by Wildschutte et al. (Wildschutte et al. 2015). Furthermore, TypeTE facilitates the genotyping with no user intervention by using as input the ‘vcf’ produced by MELT (or virtually any other pMEI detection software) to generate a new ‘vcf’ output file delivering the predicted genotypes. TypeTE also uses recently developed assemblers (SPAdes (Bankevich et al. 2012) and Minia (Chikhi and Rizk 2013) and use reads from all individuals for a locus for local MEI assembly which, in our hands, showed a higher rate of assembly than the CAP3 assembler (Huang and Madan 1999) used previously (Wildschutte et al. 2015). In addition, TypeTE can also genotype more pMEIs than previous studies based solely on *de-novo* MEI assembly (Wildschutte et al. 2015): if an incomplete Alu is assembled, TypeTE subsidize it with the exact consensus sequence based on TE’s read identity with RepBase. This additional step is performed by retrieving the subfamily on which most discordant mates align in the assembly. Here, we show that reconstruction of alternative alleles (either by local assembly or consensus-based) -- a major difference with MELT -- significantly improves the accuracy of *Alu* genotyping. Finally, TypeTE predicts the TSD accompanying each insertion and the pMEI orientation, which ensures optimal reconstruction of the two alleles. Collectively these implementations enable TypeTE to generate highly accurate *Alu* insertion genotypes.

We further tested whether the quality of the starting dataset (in particular sequencing depth) influenced TypeTE performance. By comparing results on the 1000 GP and SGDP datasets, which use different sequencing depth (on average 7.4X vs 42X, respectively), to the PCR genotypes, we found that TypeTE performs equally regardless of coverage depth (at least for reference insertions, for which we had enough loci to compare between datasets). In fact, using both non-reference and reference *Alu* insertions genotyped with TypeTE in the 1000 GP dataset, we showed that the average sequence coverage of the region flanking these loci does not seem to influence genotyping accuracy. Thus, TypeTE can support the analysis of large population dataset without stringent or highly uniform coverage requirements.

While TypeTE offers significant improvements over MELT, it still fails to genotype accurately some of the loci we experimentally assayed (16/227). Neither low sequencing coverage nor mappability issues could be readily implicated as hindering genotyping of these loci. We believe that other locus-specific idiosyncrasies prevent the ability of TypeTE to produce an accurate allele call. For instance, earlier tests on the pipeline showed that a 1-bp insertion at the end of the element in one allele or a slight error in the TSD prediction could dramatically affect the re-mapping and genotype predictions. A specific assessment of the bioinformatic methods aimed to identify TSDs should be able to improve this issue. Identifying boundaries of *Alu* insertion in low complexity (especially A-rich) regions is challenging due to inter-individual differences in the length of the poly-A tail of the element, and according to our tests, Repeatmasker often fails to identify the exact boundaries of such reference elements. Even though our pipeline in principle considers such subtle sequence variation, at least for one locus, we found that the TSD was overlapping the annotated poly-A region. Implementing changes to identify similar instances could mitigate genotyping miscalls for those loci. Additionally, our ability to evaluate the concordance of genotype predictions in low-complexity and highly repetitive regions was restrained to PCR-accessible loci. We have also noticed that altering the parameters or method for local *de novo* assembly improved the assembly of certain TE loci. An automated approach to customize the assembly parameters for each locus that failed with the standard approach would enhance the reconstruction of non-reference TE sequences. Identifying proper orientation of insertions is also crucial in accurately genotyping the insertions and we are also contemplating a read-based approach to identify the orientation of insertions in addition to the current assembly-based approach. Collecting more benchmarking data might allow us to characterize more finely these issues and adapt the pipeline accordingly. Notwithstanding these peculiar instances, TypeTE has the lowest error rate of all methods tested and as such it represents a valuable advance in the field.

The task and challenges of pMEI genotyping have been largely overlooked thus far, yet we show here that inaccurate genotyping of pMEIs can significantly bias population genetics inferences. It is presumably because of these issues that reference pMEIs have been entirely ignored in previous population genomics studies using pMEIs (L. Wang et al. 2016; L. Wang, Norris, and Jordan 2017)). By increasing genotyping accuracy for both reference and non-reference insertions, TypeTE will enhance future studies using pMEI as markers or structural variants in the human population. Notably, our results now offer a dataset of genotyped *Alu* insertions for 445 samples of the 1000 GP that is complemented by a wealth of functional data including RNA-seq (Lappalainen et al. 2013), DNA methylation (Pai et al. 2011), DNase I accessibility (Degner et al. 2012), and ATAC-seq (Kumasaka, Knights, and Gaffney 2016, 2019). We anticipate that these resources will open new avenues to explore the cis-regulatory influence of pMEIs in humans (L. Wang et al. 2016; L. Wang, Norris, and Jordan 2017; Rishishwar et al. 2018). The modularity of TypeTE allows one to easily combine new assemblers to improve the reconstruction of each pMEI, but it is also possible to skip this step and only use consensus sequence of MEI to speed up the computation time. The design of TypeTE makes it compatible with any data produced by pMEI detection tools and in principle it can be readily adapted to genotype insertions from any other retroelement families in virtually any species.

## DATA AVAILABILITY

TypeTE is freely available in the Github repository https://github.com/clemgoub/TypeTE

