## Supplementary Data for "TypeTE: a tool to genotype mobile element insertions from whole genome resequencing data"

**Supplementary Material**

**Content:**

**Figure S1.** Overview of TypeTE non-reference

**Figure S2.** Overview of TypeTE reference

**Figure S3.** Relationship between genotype concordance and mappability

**Figure S4**. Influence of the dataset of origin on variant discovery

**Figure S5.** Relationship between genotype concordance and local depth of coverage

**Table S1**. Additional PCR primer used to genotype 20 *Alu* insertion identified by the 1000 Genomes Project

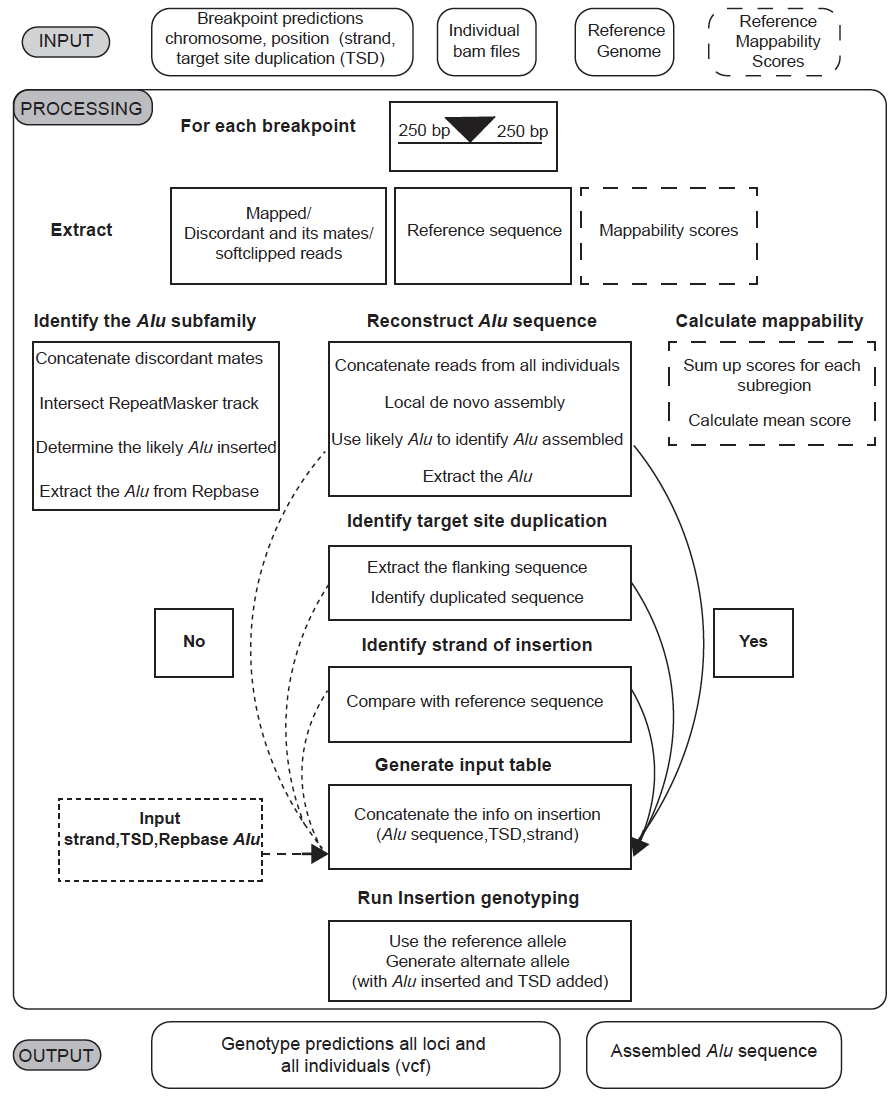

**Figure S1. Overview of TypeTE non-reference pipeline.** First, the pipeline extracts the breakpoint predictions, the strand info and target site duplications (TSDs) from the provided output of the TE discovery tool (e.g. MELT). The reference genome to which the reads are aligned in BAM files and the BAM files are needed as the input files. The reference mappability scores is an optional requirement and is used if the average mappability of the flanking region of the predicted breakpoints has to be determined. The program processes the input data by extracting the reads from the bam file from a window of +/-250 bp from each breakpoint from all individual bams provided. It also extracts the corresponding reference sequence for that window. The program also calculates the mean mappability score for that window from the reference mappability scores provided.  By intersecting the location of the mates of the discordant reads with the RepeatMasker track it determines the likely *Alu* subfamily inserted and obtains the corresponding *Alu* consensus sequence from Repbase database. The program also performs a local de novo assembly of all the reads extracted from a particular locus from all individuals. Homology based searches using the likey *Alu* consensus sequence to identify the assembled contig containing the *Alu* sequence inserted at that locus. The flanking sequence of the *Alu* from the contig is extracted and searched for TSDs. The flanking sequence is also compared with reference sequence to identify the strand of insertion. The generated information on *Alu* sequence, strand and TSDs are used to generate the input table necessary for the subsequent genotyping. The consensus *Alu* from the Repbase, TSDs and strand information provided as the input files are used whenever the pipeline failed to generate the corresponding information. The gathered information for each locus is used to generate the input table for generating reference and alternate alleles for insertion genotyping as described Figure 1A. The TypeTE outputs a ‘vcf’ containing genotype predictions for all loci for all individuals and consensus of the *Alu* sequence generated for users that are interested in doing subsequent analysis. The average mappability for flanking 250 bp regions are calculated and are included in the output vcf.

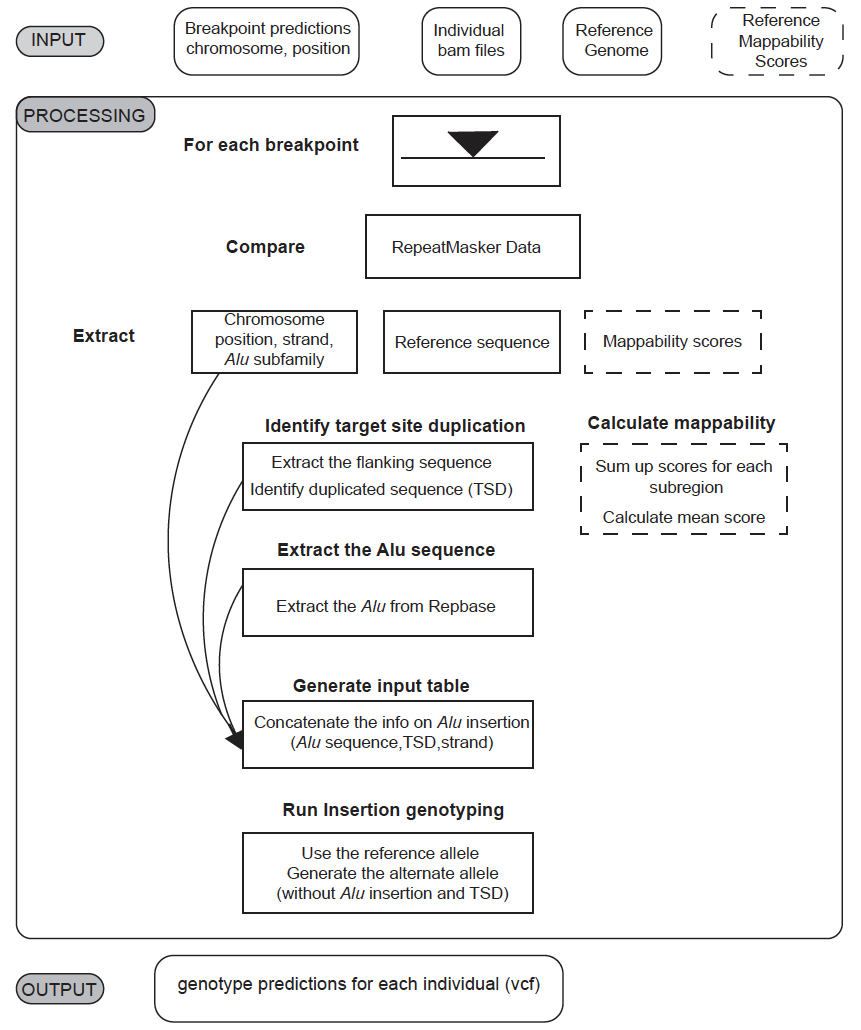
 **Figure S2. Overview of TypeTE reference pipeline.** The breakpoint predictions from a tool that detects polymorphic reference TE (e.g. MELT), reference genome to which the reads are aligned in BAM files and the BAM files are needed as the input data. The reference mappability scores is an optional requirement and is used if the average mappability of the flanking region of the *Alu* insertions has to be determined. The predicted breakpoints are compared with the RepeatMasker data to precisely identify coordinates of the *Alu* insertion, the subfamily and strand of insertion. Flanking sequences are extracted from the *Alu* coordinates for each locus to further identify target site duplications (TSDs). The program also calculates the mean mappability score for that window from the reference mappability scores provided. The gathered information for each locus is used to generate the input table for generating reference and alternate alleles for insertion genotyping as described Figure 1B. The TypeTE outputs a vcf containing genotype predictions for all loci for all individuals. The average mappability for both flanking 250 bp regions are calculated and is also included in the output vcf.

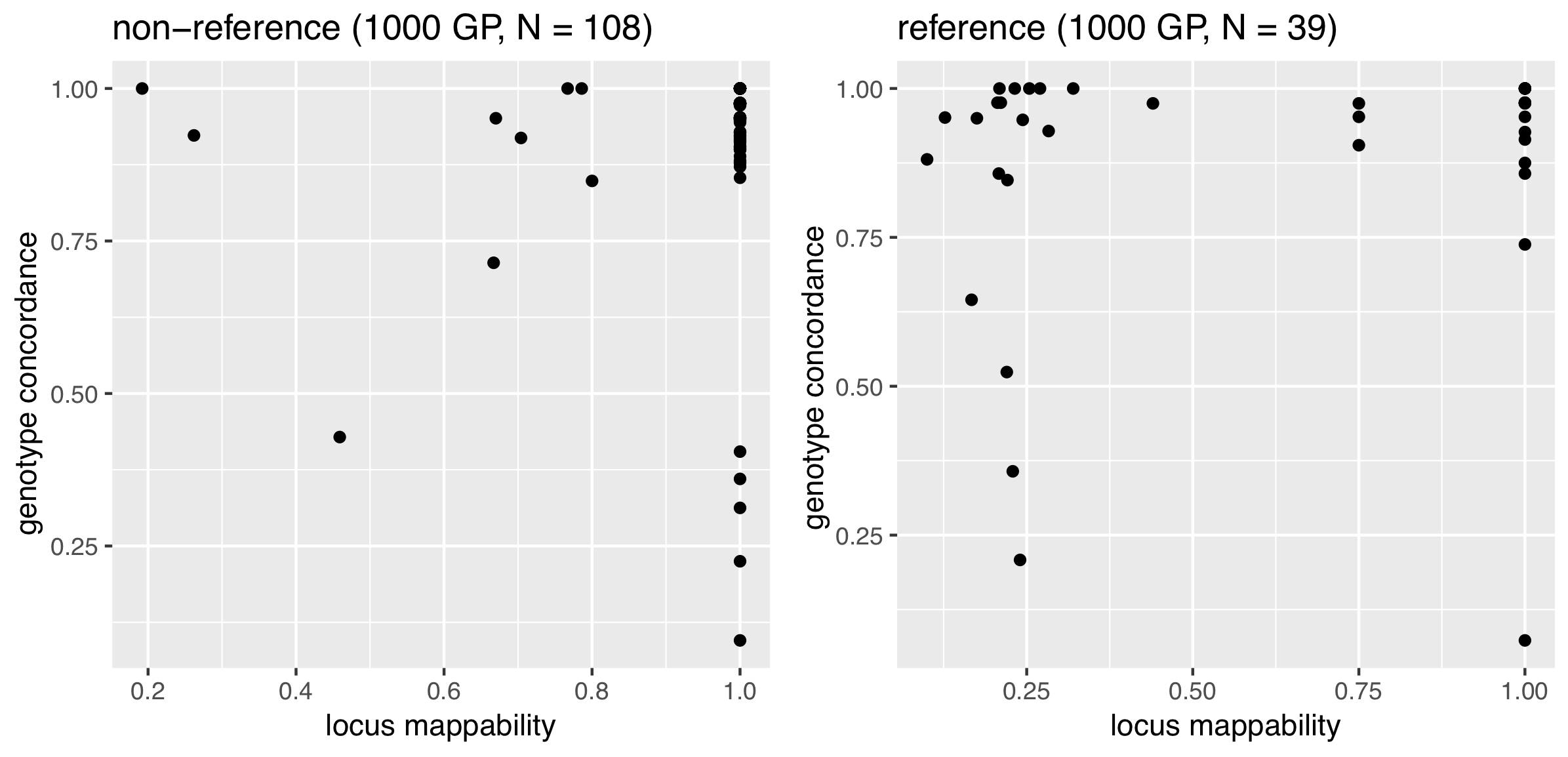
**Figure S3. Relationship between genotype concordance and mappability.** For each locus the concordance of the predicted genotype by TypeTE with the genotype obtained by PCR is plotted according to the average mappability score calculated in a +/- 500bp window of the *Alu* breakpoint.

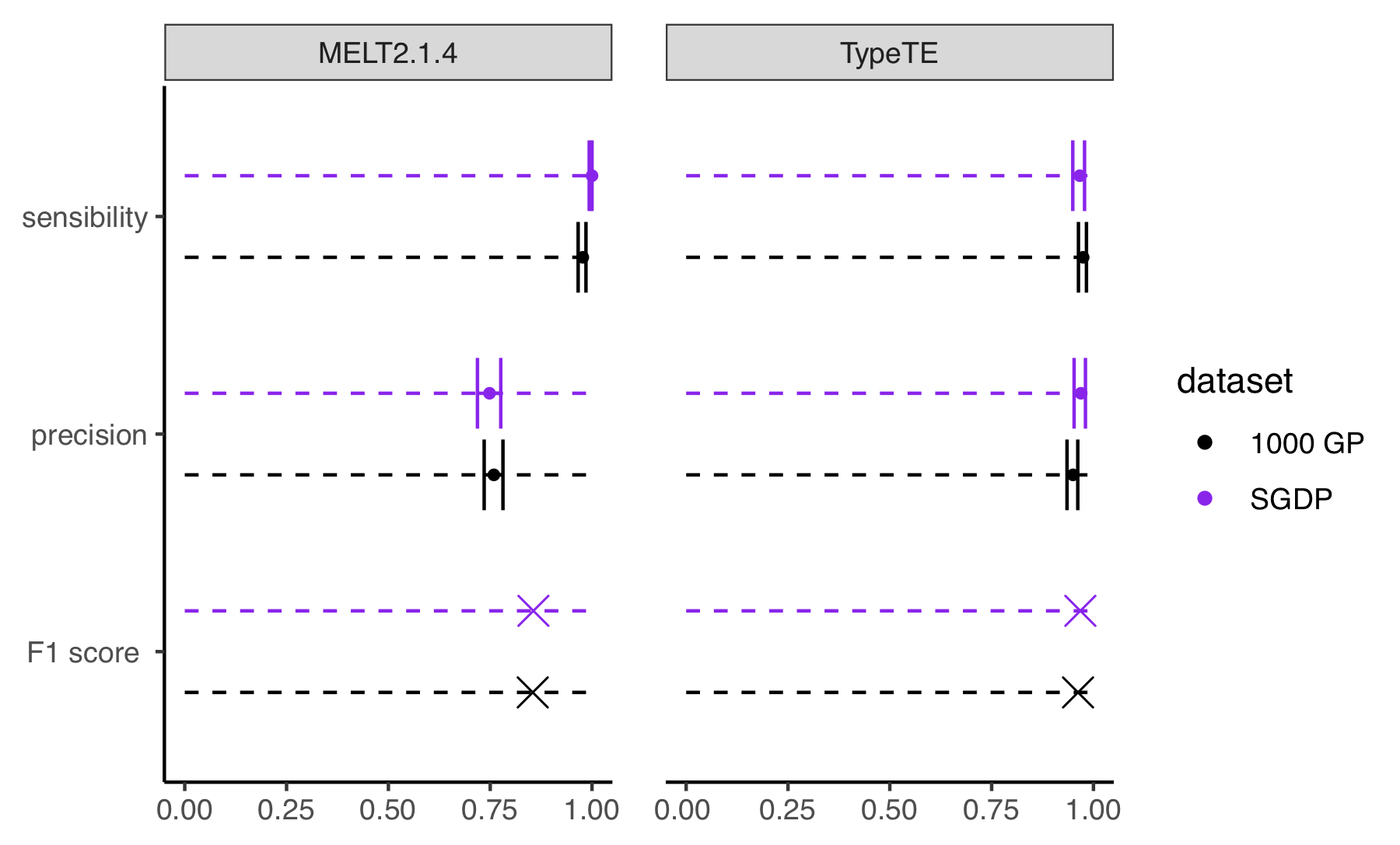

**Figure S4. Influence of the dataset of origin on variant discovery.** The sensitivity, precision and F1 scores reporting the ability of each method to accurately report a segregating site is compared for reference *Alu* between data from the 1000 GP (low coverage, black) and the SGDP (high coverage. purple).

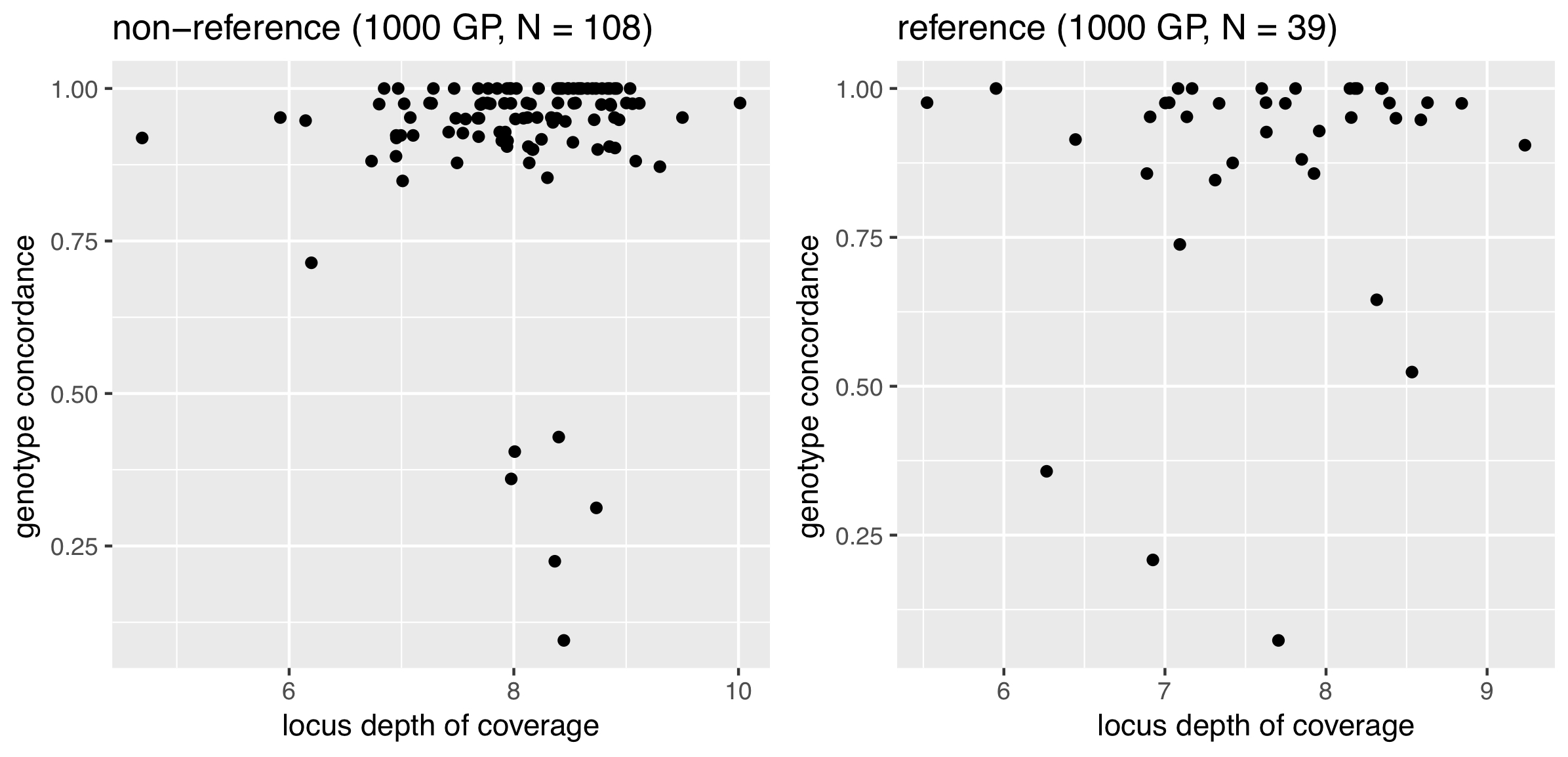

**Figure S5. Relationship between genotype concordance and local depth of coverage.** For each locus the concordance of the predicted genotype by TypeTE with the genotype obtained by PCR is plotted according to the average local depth of coverage calculated in a +/- 500bp window of the *Alu* breakpoint.

| **Burns Lab identifier** | **1KG indenfier** |  | **Forward Primer** |  | **Reverse Primer** |
| --- | --- | --- | --- | --- | --- |
| RIP-812 | DEL_pindel_37110 |  | GCTCCCTAACTTCCCTCTCTTT | | GCTCTGCACCACCTAAGGAA |
| RIP-952 | DEL_pindel_580 |  | GCCTGCAGAGTAGACAGACA |  | CTCCCCAGTGCCTCAGTAAA |
| RIP-955 | ALU_umary_ALU_277 | | CCTAGCCATGTGTCCTTCCA |  | GGCCTACCTCCTCAATTTCC |
| RIP-1024 | ALU_umary_ALU_8612 | | TGCCATGGAACTTTTCAGCC |  | AAAGATCACTGTGCTGCCTG |
| RIP-1202 | ALU_umary_ALU_11888 | | ATTGTTCTGCACGTCGGTCA |  | ATGGTTGTGAGTGGCTGGT |
| RIP-1258 | ALU_umary_ALU_12014 | | TGAGTGTGGGGAAGGTTCTC |  | GAAGAGGGAGCATCAGGACT |
| RIP-1266 | ALU_umary_ALU_12145 | | GGCTTAGGAAGGGATGGGTT |  | GTGTCTTGCAAAGGATCCGT |
| RIP-1345 | DEL_pindel_15233 |  | TCAGCAGGGTGAGAAGGATC |  | CACTTGGTTGAGTCTTGGAAATT |
| RIP-1390 | ALU_umary_ALU_5581 | | GGCTTCTCTTGTGGATGATCAG | | GCCTGGGTTCTCTGTTACCA |
| RIP-1469 | ALU_umary_ALU_7643 | | CAGGAGGAGCAGTAGGGAAG | | AGCGACAAGTTGGGTATGGT |
| 1KG_7143 | ALU_umary_ALU_7143 | | AGGCTATGGGATGGATTTGG |  | GCACCCTTGAAGTCTTGCTT |
| RIP-673A | DEL_pindel_12899 |  | CAATTCCATGAAAGCGAGGT |  | CCTCTCTCTCACAGACGGCTA |
| RIP-HL1 | ALU_umary_ALU_1992 | | TCCTGGTCTTCACATTCCCA |  | TCAGCAAAACATCCAAGGCC |
| RIP-HL2 | ALU_umary_ALU_5395 | | ATGCTGAGGGGTGTTGAAGT |  | TCCTTGTCAATAGGAGCCTCT |
| RIP-HL3 | ALU_umary_ALU_10819 | | GCAATGCATGAGTGTACATATGT | | GTCAGGCCTGCTCACTATAGT |
| RIP-HL4 | DEL_pindel_31897 |  | ACTGTCTGTCTTGGGTTATTTCT | | CATGACAGGACAAGGGGAGT |
| RIP-HL5 | DEL_pindel_32884 |  | GGAGTGAGGGCTATTGGACA |  | ATCTACGTAGCCTTTCTCTTTGT |
| RIP-HL6 | DEL_pindel_6469 |  | TGTTAGGTCAAAAGACAGGCAC | | TGGCTTAGCACACTTAAGAATCA |
| RIP-HL8 | ALU_umary_ALU_8297 | | TCAGGTGAGATTGAGGTCTGT | | ACAAGATCCCTTTGGCAAACA |
| RIP-HL9 | ALU_umary_ALU_5043 | | CTGAATTGCAAAACTCTTCCCC | | GGAACATGACCTCGGATTGC |

**Table S1. Additional PCR primer used to genotype 20 *Alu* insertion identified by the 1000 Genomes Project.**
